# Enhancing the dark side: Asymmetric gain of cone photoreceptors underpins discrimination of visual scenes based on their skewness

**DOI:** 10.1101/2021.04.29.441966

**Authors:** Matthew Yedutenko, Marcus H.C. Howlett, Maarten Kamermans

## Abstract

Psychophysical data indicates humans can discriminate visual scenes based on their skewness – the ratio of dark and bright patches within a visual scene. It was also shown that on a phenomenological level this skew discrimination is described by the so-called Blackshot mechanism, which accentuates strong negative contrasts within a scene. Here we demonstrate that the underlying computation starts as early as the cone phototransduction cascade whose gain is higher for strong negative contrasts than for strong positive contrasts. We recorded from goldfish cone photoreceptors and found that the asymmetry in the phototransduction gain leads to higher amplitude of the responses to negatively than to positively skewed light stimuli. This asymmetry in the amplitude was present in the photocurrent, voltage response and cone synaptic output. These results highlight the importance of the early photoreceptor non-linearity for perception. Additionally, we found that stimulus skewness leads to a subtle change in photoreceptor kinetics. For negatively skewed stimuli, the cone’s impulse response functions peak later than for positively skewed stimulus. However, stimulus skewness does not affect the cone’s overall integration time.

## Introduction

Psychophysical studies show that humans are sensitive to the ratio of negative (intensity lower than the mean) and positive (intensity higher than the mean) patches of contrast in visual scenes (Chubb *et al*., 1994, 2004; Graham *et al*., 2016). This ratio is described by the parameter known as skewness. Visual stimuli are called positively skewed if there is a predominance of negative contrasts with some infrequent patches of high positive contrast and are called negatively skewed when the situation is reversed. Figure 1 illustrates one’s ability to discriminate visual scenes based on skewness by mimicking an experiment performed by Chubb et al. ( 1994, 2004). The textures were randomly drawn from two distributions equal in every aspect but skewness (Bonin *et al*., 2006). Yet, one can appreciate the clear difference between negatively skewed images (right upper) and positively skewed images (remaining panels). Chubb et al. (1994, 2004) showed that on a phenomenological level the sensitivity to skewness can be described by the so-called Blackshot mechanism. This Blackshot mechanism does not react to skewness per se, rather its sensitivity to strong negative contrasts is simply much higher than to strong positive contrasts and so it effectively reports the fraction of strong negative contrasts within the scene.

**Figure 1.**
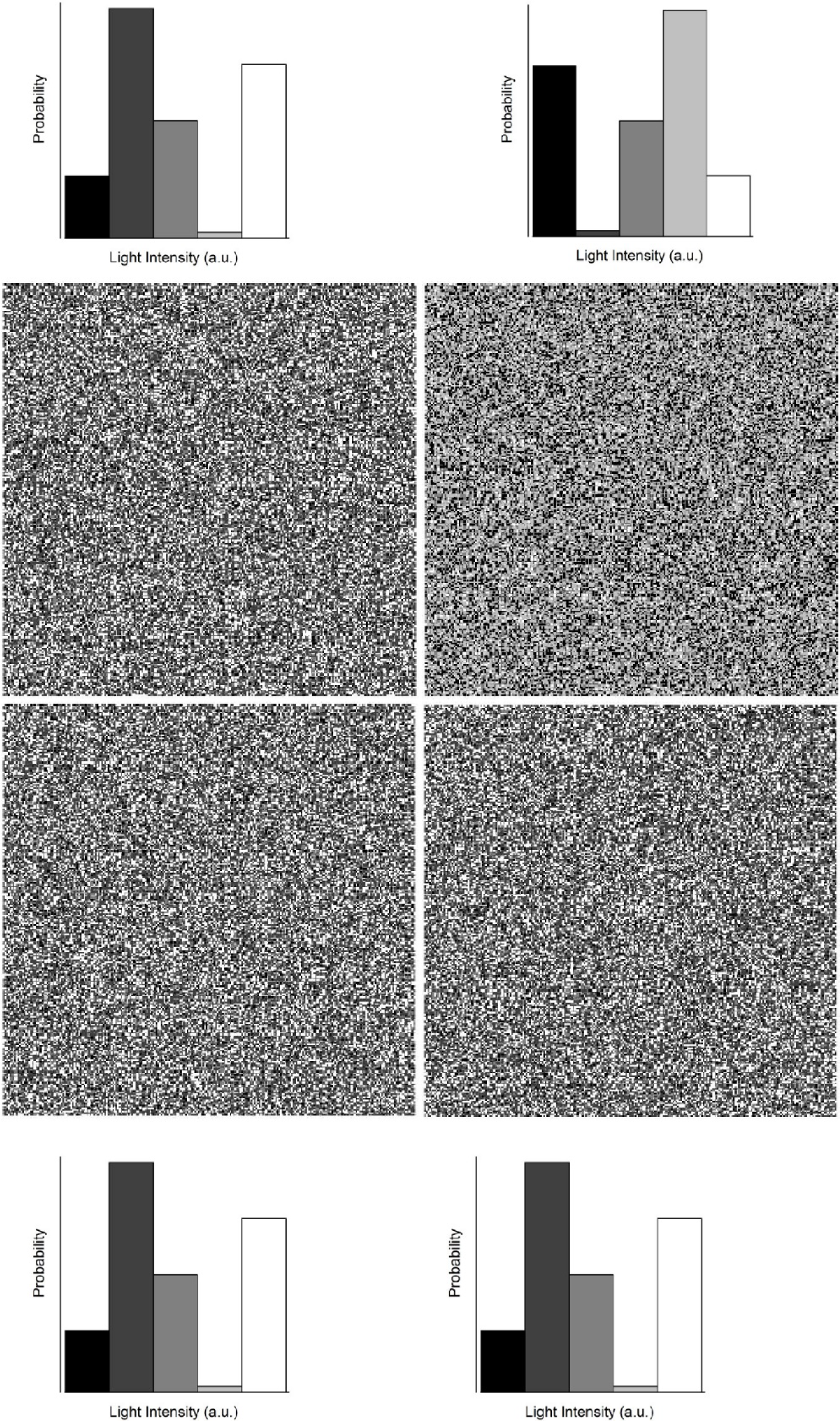
The discrimination of skewed stimuli. An example of textures used by Chubb et al.( 1994, 2004) to psychophysically probe the ability of humans to discriminate visual scenes based on their skewness. The textures were randomly drawn from the probability distributions adjacent to each panel following the approach described by Bonin et al. (2006) and differed only in terms of skewness. The upper right texture is negatively skewed (-0.4), the remainder are positively skewed (+0.4).

What are the neuronal correlates of the Blackshot mechanism? Studies using salamander retinal ganglion cells (RGC) (Tkačik *et al*., 2014) and cat lateral geniculate nucleus neurons (LGN) (Bonin *et al*., 2006) did not report any response differences associated with changes in stimulus skewness. Therefore, both studies concluded that the discrimination between skewed stimuli occurs in the visual cortex. On the other hand, it is well-established that the retinal photoreceptor’s gain is asymmetric: given an equal input magnitude, the response amplitude to a strong (>0.4 Weber unit) negative contrast step is greater than it is to a strong positive contrast step (Laughlin, 1981; van Hateren, 2005; Endeman & Kamermans, 2010; Baden *et al*., 2013; Angueyra *et al*., 2021). Furthermore, this responses asymmetry is observed throughout the post-receptor retinal stages (Lee *et al*., 2003; Zaghloul *et al*., 2003), the LGN (Kremkow *et al*., 2014), and the primary visual cortex (Zemon *et al*., 1988; Jin *et al*., 2008; Yeh *et al*., 2009; Kremkow *et al*., 2014). The asymmetrical processing of positive and negative contrasts should lead to different response amplitudes to negatively and positively skewed stimuli and thus might underpin the discrimination of skewed stimuli.

A possible reason why the differences in responses to skewed stimuli were not found in RGC (Tkačik *et al*., 2014) and LGN (Bonin *et al*., 2006) studies was the power spectra of the stimuli used. In both cases, the researchers employed band-limited white noise, where large proportions of the signal power are outside the photoreceptor frequency bandwidth. Thus, in both studies temporal filtering discarded a significant portion of the signal, reducing the skewness and amplitude of the “effective” light stimuli actually available to photoreceptors. Consequently, Bonin et al. ( 2006) and Tkacik et al. ( 2014) may not have found significant differences in the processing of skewed stimuli because a) their stimuli hardly differed in terms of “effective” skewness and b) the “effective” amplitudes of the employed stimuli were too low to drive cone photoreceptors outside their linear response range.

Here we address whether photoreceptors process positive and negative skewed stimuli differently. To account for the kinetics of cone photoreceptors, we stimulated goldfish cones with sets of skewed stimuli with bandwidth similar to those of the goldfish cones. We found the asymmetry in goldfish photoreceptor gain does indeed entail skew-dependent changes in the cone response, leading to greater response amplitudes to negatively than to positively skewed stimuli. This asymmetry originates in the cone phototransduction cascade and is the first step in implementing the Blackshot mechanism. Additionally, we found that the cone’s impulse response function peaks ≈ 3.6 ms later for negatively skewed stimuli whereas the cone’s integration time is unaffected by stimulus skewness.

Thus, we show that in contrast to what models of various retinal, thalamic and cortical neurons (Carandini *et al*., 2005; Schreyer & Gollisch, 2021) often imply, cone photoreceptors do not simply linearly relay visual stimuli to downstream circuitry. Rather, they also emphasize specific features of their stimuli. These results highlight the important of accounting for the full non-linear character of photoreceptor signal processing when studying the visual system.

## Materials and Methods

### Recording procedures

All animal experiments were conducted under the responsibility of the ethical committee of the Royal Netherlands Academy of Arts and Sciences (KNAW), acting in accordance with the European Communities Council Directive of 22 July 2003 (2003/65/CE) under license number AVD-801002016517, issued by the Central Comity Animal Experiments of the Netherlands. In all experiments retinas of adult goldfish (*Carassius auratus*) were used.

Goldfish were first dark-adapted for 5-10 minutes, sacrificed and the eyes enucleated. Retinas were isolated under dim red illumination then placed photoreceptor side up in a recording chamber (300 μl, model RC-26G, Warner Instruments) mounted on a Nikon Eclipse 600FN microscope. The preparation was viewed on an LCD monitor by means of a 60× water-immersion objective (N.A. 1.0, Nikon), a CCD camera, and infrared (λ > 800 nm; Kodak wratten filter 87c, United States) differential interference contrast optics. Tissue was continuously superfused with oxygenated Ringer’s solution at room temperature (20°C). The composition of Ringer’s solution was (in mM): 102.0 NaCl, 2.6 KCl, 1.0 MgCl_2_, 1.0 CaCl_2_, 28.0 NaHCO_3_, 5.0 glucose continuously gassed with 2.5% CO_2_ and 97.5% O_2_ to yield a pH of 7.8 (osmolarity 245–255 mOsm). For calcium current (I_Ca_) measurements, 5 mM of NaCl was substituted with 5mM of CsCl and 100 μM of niflumic acid added.

Measurements from goldfish cones were performed in current-(voltage response), and voltage-(photocurrent), clamp configurations. Patch pipettes (resistance 7-8 MOhm, PG-150T-10; Harvard Apparatus, Holliston, Massachusetts) were pulled with a Brown Flaming Puller (Model P-87; Sutter Instruments Company). Patch pipette solution contained (in mM): 96 K-gluconate, 10 KCl, 1 MgCl2, 0.1 CaCl2, 5 EGTA, 5 HEPES, 5 ATP-K2, 1 GTP-Na3, 0.1 cGMP-Na, 20 phosphocreatine-Na2, and 50 units ml^−1^ creatine phosphokinase, adjusted with KOH to pH 7.27–7.3 (osmolarity 265–275 mOsm). The chloride equilibrium potential (E_Cl_) was -55mV except when the calcium current (I_Ca_) was studied. Here E_Cl_ was set at -41 mV by changing the concentrations of K-gluconate and KCl to 87 and 19 mM, respectively. All chemicals were supplied by Sigma-Aldrich (Zwijndrecht, the Netherlands), except for NaCl (Merck Millipore, Amsterdam, the Netherlands).

Filled patch pipettes were mounted on a MP-85 Huxley/Wall-type manual micromanipulator (Sutter Instrument Company) and connected to a HEKA EPC-10 Dual Patch Clamp amplifier (HEKA Elektronik GmbH, Lambrecht, Germany). After obtaining a whole cell configuration, cones were first spectrally classified then stimulated with a set of skewed stimuli of corresponding chromaticity. Data were recorded at a sample rate of 1 kHz using Patchmaster software package (HEKA Elektronik GmbH). In total, we recorded 14 cones in voltage-clamp mode (8 light responses, 6 measurements of I_Ca_) and 16 cones in current-clamp mode (all light responses). The liquid junction potential (approximately -15 to -16 mV) was not compensated.

### Light stimuli

The light stimulator was a custom-built LED stimulator with a three-wavelength high-intensity LED (Atlas, Lamina Ceramics, Westhampton, New Jersey, US). The peak wavelengths were 465, 525 and 624 nm. Bandwidth was smaller than 25 nm. Linearity was ensured by an optical feedback loop. The output of the LED stimulator was coupled to the microscope via an optic fiber and focused on cone outer segments though a 60× water-immersion objective. The mean light intensity of all stimuli was 1.2*10^4^ photons/µm^2^/s, which is in the photopic level for goldfish (Malchow & Yazulla, 1986). Stimuli were presented at 1 kHz.

#### Skew Stimulus Set#1

Skewed stimuli were based on the natural time series of chromatic intensities (NTSCI) from the Van Hateren library (Van Hateren *et al*., 2002). Given that psychophysical studies have shown visual scene discrimination occurs when intensities within the spatial domain are skewed (Figure 1), one may wonder how appropriate it is to study the phenomena with time series of intensities. However, we argue such an approach is correct for the following two reasons. Firstly, saccadic eye movements convert spatial stimuli into time series. Hence, the photoreceptors of subjects in psychophysical studies ‘perceive’ the presented skewed textures as time series of intensities. Secondly, a photoreceptor’s direct light response only depends on the light intensities falling upon its outer segment and not on the stimulus’s spatial structure. Hence, one can reproduce the response of an array of cones to the presentation of a single spatial texture by recording the response of a single cone to a collection of time series of intensities that correspond to the differing retinal locations.

The NTSCI power spectra is typical of that of ‘natural stimuli’ in that power declines as a function of frequency(Van Hateren, 1997; Van Hateren *et al*., 2002; Frazor & Geisler, 2006). As a result of the predominance of lower frequencies, most of the light intensity changes throughout the NTSCI occur over timescales accessible to goldfish cones and previously the NTSCI has been used to unlock several non-linear performance features of cones (Endeman & Kamermans, 2010; Howlett *et al*., 2017). To ensure that all aspects other than skewness remained equal, we first picked short stretches from the NTSCI that were positively skewed, then simply flipped these around the mean to generate negatively skewed stimuli.

To generate Stimulus set#1 we divided the NTSCI (Van Hateren *et al*., 2002) into one-second long stretches. Then from each stretch we subtracted its minimum value, adjusted their mean light intensities to be equal and picked stretches with similar power spectra and root mean square (r.m.s.), and median contrasts (between 0.23 and 0.25). The r.m.s. and median contrast for each stretch was calculated respectively as the ratio between the stretch’s standard deviation and its mean, and the ratio between its deviation from the median, and its median. To ensure an absolute similarity between positively and negatively skewed stimuli, we selected only stretches where the maximum value was not larger than 2 times the mean. Next, we chose stretches with skewness’s of 0.9, 1.6, 2.2. The skewness was calculated with the equation (1):

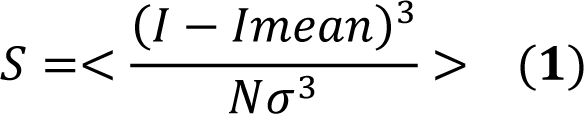

where N is the number of elements in the stretch, I corresponds to the light intensity of an element, I_mean_ and σ are mean and standard deviation within the stretch and brackets denotes averaging over the period.

We further narrowed our selection to three stretches all with similar power spectra (data not shown). Power spectra were calculated by Welch’s averaged periodogram method (Welch, 1967). No window function was used, the length of the Fourier transform was same as the length of each corresponding data sequence. Total stimulus power was calculated as the integral under power spectra, the differences in the total stimulus power were no more than 10%. Finally, an additional pink noise stimulus with zero-skew and similar power spectra was added to the set. In total, Skew Stimulus set#1 consisted of seven 1-second stimuli.

#### Skew Stimulus set#2

This stimulus set consisted of three 4-second long stretches with a skewness of 2.2, 0, and -2.2. They were generated in the same way as the Skew Stimulus set#1, but with one additional condition: the degree of skewness delivered by the stimulus remained unchanged by the cone’s temporal filtering. This was ensured by first convolving the NTSCI stretch with the mean photocurrent impulse response function (see below) obtained from responses to Skew Stimulus set #1. The skew of the convolution product, representing the “effective” stimulus, was then compared with the skew of the original stimulus (Figure 3). This was further confirmed by determining the effective skewness after convolving the stimuli with the impulse response function of each cone measured under Skew Stimulus set #2 conditions (Figure 3B).

### Calcium current isolation

To measure I_Ca_, we used the mean voltage response (7 cells, 69 repeats in total) of cone photoreceptors to Stimulus set #2 as the command voltages for the voltage clamp experiments.

To isolate I_Ca_ we followed the approach described by Fahrenhoft et al. (Fahrenfort *et al*., 1999). Briefly, to eliminate the calcium-dependent chloride current E_Cl_ was set at -41 mV and 100 µM of niflumic acid added to the Ringer’s solution; delayed rectifying, and hyperpolarization-activated, potassium currents were blocked by substituting 5 mM of NaCl in Ringer’s solution with 5 mM of CsCl; light-activated conductances were saturated by a 20 µm spot of bright white light focused on the cone outer segment; linear leak currents were removed by subtraction. The leak current was estimated by clamping cones at -70mV, stepping to potentials between -100 and 20 mV in 5 mV steps for 100 ms, calculating the mean current between 20 and 60 ms after the step onset, then determining the linear fit of the IV-relation between -100 and -60 mV (Vroman *et al*., 2014; Kamar *et al*., 2019).

### Data analysis

For each cell, the skewness of its mean response to each stimulus was determined using equation (1). In Figures 2, 3D and 8A, data was fitted using build-in Matlab least square methods. All data analysis was performed in Matlab and Python.

**Figure 2.**
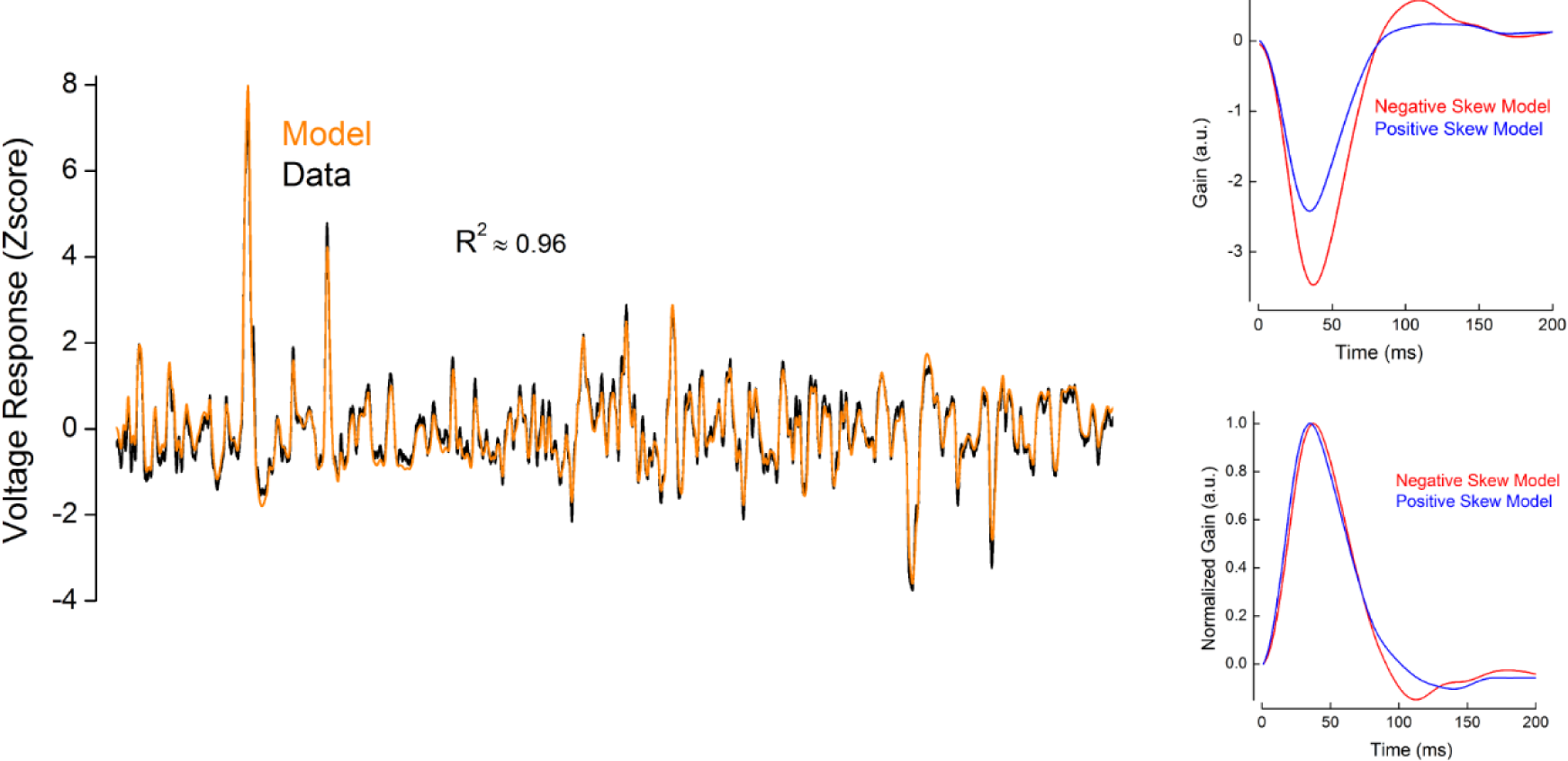
Example of the performance of the Van Hateren model for goldfish cones. Left. Voltage responses as Z-scores to Skew Stimulus set #2 for a representative recorded cone (black line) and for the simulated cone (orange line). The coefficient of determination (R^2^) between these two traces was 0.98. Right. Impulse response functions obtained from the simulated responses to negatively (red) and positively (blue) skewed stimuli. Parameters for the simulation are listed in the Table 1.

**Table 1.**
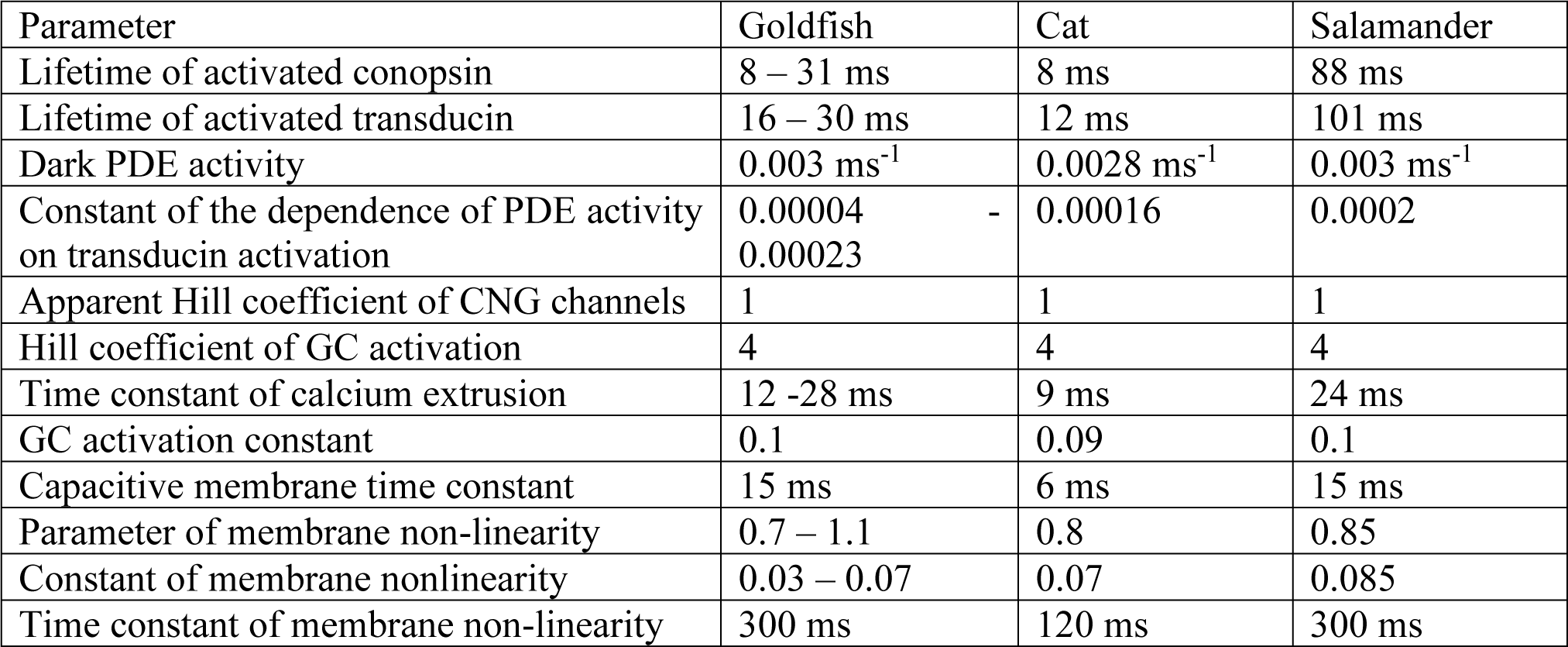
Parameters of the Van Hateren model used to simulate the responses of goldfish, salamander and cat cones. For the goldfish, model parameters were obtained by fitting cones responses (n=7) to Skew Stimulus Set #2 while constraining the range the parameters varied to within that determined by Endeman and Kamermans (2010). For the cat and salamander, parameters were adjusted such that the simulated-cone’s impulse response function time-to-peak approximately matched that estimated, respectively, by Donner and Hemila (1996) and by Rieke (2001) and Baccus and Meister (2002).

Parallel cascade identification is the most rigorous method to describe the signal processing properties of cone responses to naturalistic stimuli (Korenberg, 1991) . However, for practical reasons our analysis only focuses on the estimation of the first parallel cascade, which is effectively a linear filter followed by a static non-linearity. Apart from the computational and descriptive simplicity, this approach is also justifiable as it accurately describes cone responses, accounting for over 95% of the variance (Figure 8A).

The linear temporal filtering properties of a cone was described by its impulse response function. Impulse response functions were estimated as the inverse Fourier transform of the ratio between stimulus-response cross-power and stimulus power spectrum (Wiener, 1964; Kim & Rieke, 2001). The spectra were calculated using Welch averaged periodogram method (Welch, 1967). Stimuli and responses were detrended, divided into 500 ms long stretches, with 50% overlap, and windowed with a hamming function. The length of the Fourier transform was 1024 ms. Cone integration time was calculated as the integral of the impulse response function’s initial polarization lobe (Daly & Normann, 1985).

To estimate the effective” stimuli perceived by each cone photoreceptors during each stimulus, we convolved the cone’s impulse response function during that condition with the original light stimulus. Skews of these “effective” stimuli were calculated with equation (1). Discrepancies between the skewness of the “effective” stimuli and the skewness of the cone’s responses, were considered a result of non-linear cone properties.

For “effective” Weber contrast steps (Figures 7&8A), light stimuli were first converted into Weber contrasts steps (Figure 7) with equation (2):

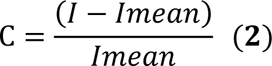

These Weber contrast steps were then convolved with a mean impulse response function to obtain the “effective’’ Weber contrast steps. The mean impulse response function used here was the averaged voltage-response derived impulse response function of all 16 cones measured in current clamp, which was subsequently scaled such that the integral under its curve yielded one (Figure 7; Howlett et al., 2017). Even though a cone’s impulse response function is affected by the skewness of the stimulus (Figure 8B&C), we used this mean impulse response function for the following reasons.

Firstly, observed changes to the impulse response function shape had little effect on the cone response dynamics. We convolved Skew Stimulus Set #2 with impulse response functions derived from responses to positive and negative stimulus of each of the cones (n=7) and found that the resulting convolution products were highly correlated (r≈0.98 ± 0.01). Secondly, averaging across different stimulus skew conditions was crucial to account for biases in the estimate of the amplitude of the impulse response function arising from skewness of the light stimuli (Chichilnisky, 2001; Simoncelli *et al*., 2004; Bonin *et al*., 2006; Tkačik *et al*., 2014). Finally, the goal of this analysis was to illustrate cone photoreceptor gain asymmetries rather than to provide a veridical description of the gain dependence on stimulus contrast.

To estimate the cone’s non-linear gain function parameters we fitted the relationship between all the mean cone voltage responses and the “effective” Weber contrast steps (Figure 8A, 19000 data points) as the power function of input contrast (Van Hateren & Snippe, 2006) with equation 3:

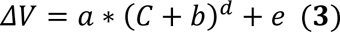

Here ΔV denotes voltage response, C is the “effective” Weber contrast, a, b, d, and e are fit parameters. At the biophysical level b) corresponds to the baseline rate of phosphodiesterase activity (PDE), d) describes the inter-dependence between the PDE activity rate and the voltage response, e) is proportional to the baseline concentration of cyclic guanosine monophosphate and a) is a scaling factor (Van Hateren & Snippe, 2006). The quality of the fit was quantified with the adjusted coefficient of determination (R^2^). The highest R^2^ (0.95) was obtained using the following parameter values (value ± 95 % confidence interval): a=0.05138 ± 0.03506, b=1.166 ± 0.072, d= -0.1251 ± 0.0959, e=-0.05046 ± 0.03552. Our estimation of the power function (d) was close to that obtained by Van Hateren and Snippe (2006) in their theoretical study.

### Model

Photoreceptor responses were modelled in matlab using Van Hateren’s model of vertebrate photoreceptors (van Hateren & Snippe, 2007), which was shown to be remarkably precise in capturing the processing steps involved in generating a cone’s signal. Apart from the activation of hyperpolarization-activated current (I_h_; Howlett et al., 2017; Kamermans et al., 2017), the model closely simulates all the cone’s biophysical processing steps from the photon-initiated activation of conopsins to the cGMP-regulated changes in the photocurrent, followed by the generation of the voltage response. The model simulates cone photoreceptors as a cascade of low-pass filters, a static (instantaneous and memoryless) non-linearity, and two divisive feedback loops (van Hateren, 2005; van Hateren & Snippe, 2007). The low-pass filters correspond to the kinetics of the different biophysical processing steps. The non-linearity describes the inverse proportional dependence between light intensity and changes in the cGMP concentration. The first feedback loop describes the regulation of the rate of cGMP production by calcium influx through cGMP-gated channels. The second feedback loop corresponds to the regulation of the membrane voltage by voltage-sensitive channels in the cone inner segment. The cone’s non-linear gain (Figure 8A) originates from the interplay between the hydrolysis of the cGMP by PDE and calcium-regulated (feedback loop) production of the cGMP by guanylyl cyclase (GC)

We verified that the Van Hateren model could capture responses to skewed stimuli. For this we fitted the model to the voltage responses of 7 goldfish cones recorded under Skew Stimulus Set#2 conditions. The model parameters were modulated within the ranges determined by Endeman and Kamermans (2010) and are shown in the Table 1. For all 7 cells, the correlation coefficient between modelled and recorded voltage responses were no less than 0.97 (adjusted R^2^ ≥ 0.94 Figure 2). Moreover, the impulse response functions estimated from the simulated responses to positively and negatively skewed stimuli retained features of the impulse response functions derived from the recorded voltage responses. For example, for both recorded and simulated cone voltage responses, impulse response functions peaked 3 ms (8.5±1%, p=0.00013) later under the negatively skewed stimulus compared to the positively skewed condition but showed no statistically significant difference in their full width at half maximum (FWHM), or in integration time (Figure 2). Thus, Van Hateren’s model reproduces accurately cone responses to skewed stimuli.

Next, we used the Van Hateren model to estimate the “effective” stimuli perceived by salamander and cat cones in the studies by Tkacik et al. (2014) and Bonin et al. (2006), respectively. To model salamander cones we adjusted the parameters of Van Hateren’s model such that the time course of the impulse response functions of simulated cones resembled those of salamander cones reported by Rieke (2001) and Baccus and Meister (2002). The exact simulation parameters are reported in the Table 1. Similarly, to model cat cones the Van Hateren model parameters were adjusted so that the impulse response function time course resembled the estimates made by Donner and Hemila (1996). For the cat, exact parameters of the simulation are reported in the Table 1.

“Light” stimuli mimicking those used by Bonin et al. and Tkacik et al. (2014) where respectively used to study the responses of the modelled cat (Figure 9) and salamander (Figure 10) cones to changes in skewness. The only difference was that for illustrative ease the positively and negatively skewed stimuli were mirror copies of each other. Cat stimuli had a r.m.s. contrast of 0.7, skews of ±0.4 and a flat power spectrum bandlimited to 124 Hz. Salamander stimuli had a r.m.s. contrast of 0.2, skews of ±2 and a flat power spectrum bandlimited to 30 Hz.

### Statistics

All data are presented as mean ± SEM unless otherwise stated. Differences between groups were tested using two-tailed t-test (Student, 1908).

## Results

### Cone responses vary with skewness

Direct light responses of photoreceptors are not affected by the spatial structure of stimuli and instead only depend on the intensity of light falling on the photoreceptor outer segment (see Material and Methods-Light stimuli). Hence, to assess the photoreceptor’s contribution to skew discrimination, we exposed goldfish cones to a series of modified naturalistic time series of chromatic intensities (NTSCI) from the Van Hateren library (Van Hateren et al., 2002; Skew Stimulus set#1 in Material and Methods) and recorded their photocurrent and voltage responses (Figure 3A). Stimuli were equal in terms of mean intensity, root mean square contrast and median contrast, and had similar power spectra, while their skewness varied from -2.2 to +2.2. Positively and negatively skewed stimuli were mirror copies of each other, therefore any asymmetries between corresponding responses would reflect an asymmetry in the cone’s processing.

**Figure 3.**
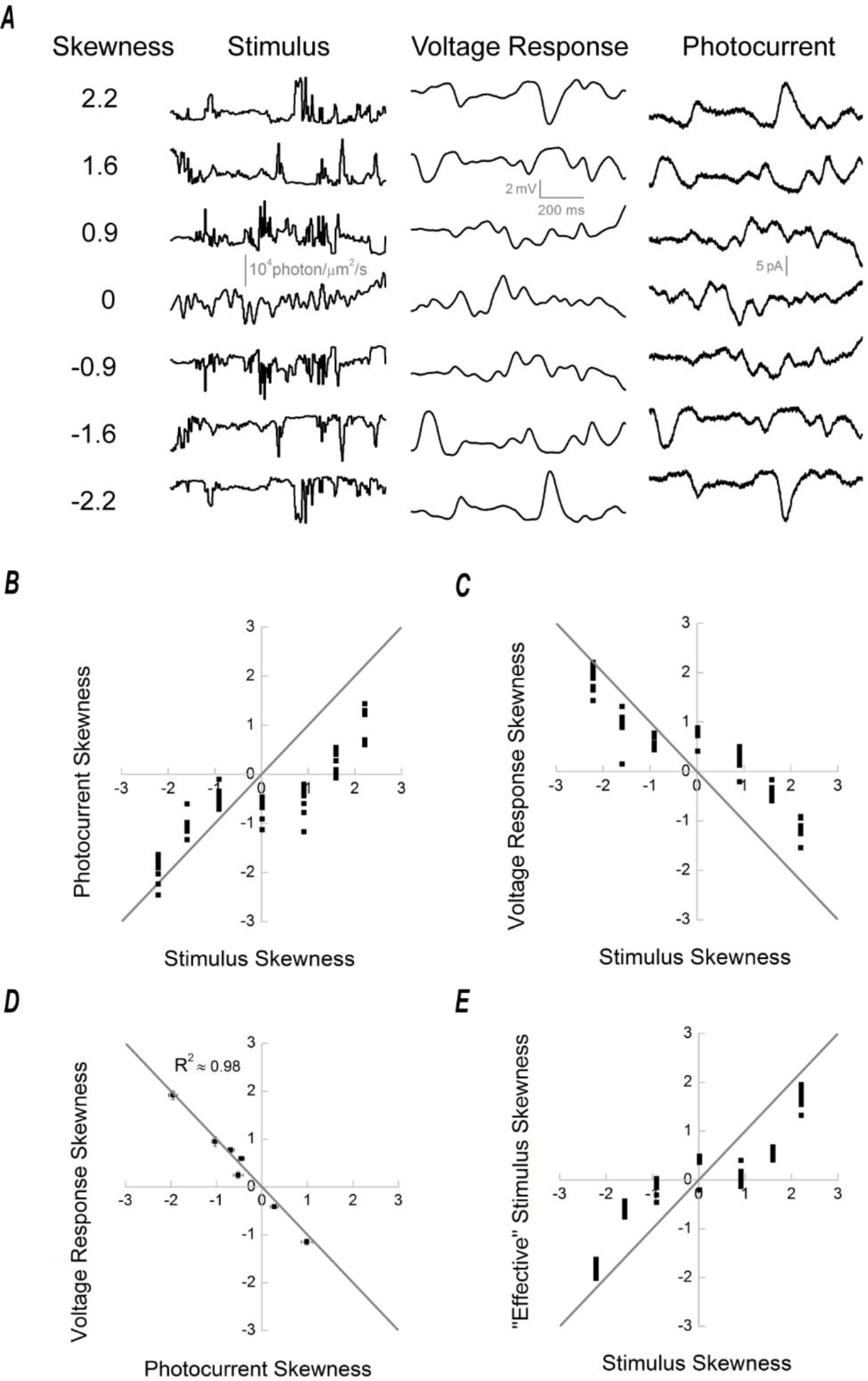
Cone processing affects stimulus skewness. **A.** Skew Stimulus set#1 (left) together with examples of voltage (middle), and photocurrent (right), responses recorded from cone photoreceptors. **B.** Photocurrent skewness as a function of stimulus skewness for Skew Stimulus set #1 (n=8). Each black square represents the skewness of a cell’s photocurrent response to a skewed stimulus condition. The grey line depicts the situation where the response skewness is equal to the stimulus skewness. The panel shows cone responses to positive skewed stimuli were less skewed than their stimulus, but to stimuli with zero or negative skews cone responses were more skewed than their stimulus. **C.** Voltage response skewness as a function of stimulus skewness (n=9). Note that the voltage responses and stimuli skews have different sign due to the signal sign-inversion. The grey line describes the situation where response and stimulus skews are equal in magnitude. As was the case in B), cone responses to positive skewed stimuli were less skewed than their stimulus. To stimuli with zero or negative skews, the skewness of the cone’s response was greater than that of its stimulus. **D.** Skewness of the voltage response as a function of the photocurrent. For each stimuli condition, the skews of each corresponding voltage response, and photocurrent, were averaged and the resulting means (±SEM) plotted as a voltage response verses photocurrent function. Note that as the cone’s voltage response is sign-inverted relative to its photocurrent their skews are also sign inverted. The situation where the voltage response and photocurrent have equal magnitude of skewness is described by the grey line, which fits the data with adjusted R^2^ of ≈ 0.98. This indicates that the asymmetric processing of positively and negatively skewed inputs originates in the phototransduction cascade. **E.** “Effective” skewness perceived by the cones as the function of the skewness of the light stimuli. “Effective” skews were estimated from the convolution product of the light stimuli with the cone’s impulse response function (Figures 4A&B). The grey line depicts the situation, where “effective” and response skews are equal. Note that for illustrative convenience the “effective” skewness estimated from voltage responses were multiplied by -1. Figure 3E indicates that even though naturalistic stimuli were used, some aspects of the stimuli were still unavailable to drive cone responses on account of the cone’s temporal filtering properties. This in turn reduced the range of “effective” skews by almost 30% relative to the original -2.2 to +2.2 range of stimulus skews.

To determine whether cones process negatively and positively skewed light stimuli differently, we plotted the skews of the photocurrent (Figure 3B) and voltage responses (Figure 3C) against the skews of the light stimuli. If there is no difference in processing, the skewness of the response will be equal to the light stimulus skewness and thus the data points will fall along a straight slope. However, if there is an asymmetry in the processing of positive and negative contrasts it would necessarily lead to the deviation of the data points from the grey line. Figures 3B&C shows that for positively skewed stimuli, the photocurrent and voltage responses are skewed to a lesser degree than are the light stimuli, whereas for the negatively skewed stimuli they are almost as equally skewed as the light stimuli. Note that the signal sign-inversion of the voltage response also sign-inverts its skewness. Figures 3B&C indicate an asymmetry in the processing of negatively and positively skewed stimuli by cone photoreceptors.

### The processing asymmetry originates exclusively within the phototransduction cascade

What are the cellular mechanisms leading to the differences in the processing of negatively and positively skewed stimuli? To tease apart the relative contributions of the phototransduction cascade and the voltage activated membrane conductances, we plotted the skewness of the voltage responses and photocurrent against each other in Figure 3D. The grey line depicts the condition where photocurrent and voltage response skews are equal in magnitude. All data points fall on this line (R^2^ =0.98), meaning that the skewness of the photocurrent accounts for 98% of the skewness of the voltage responses. This means that the phototransduction cascade is the primary source of the asymmetric processing of the positively and negatively skewed stimuli.

### Temporal filtering affects stimulus skewness

A cone’s finite kinetics may act to temporally filter our light stimulus and thus affect the skewness of the “effective” stimulus perceived by a cone. To determine the “effective” stimuli skews we first estimated the temporal filters of the cone’s photocurrent responses to each of the skewed stimuli stretches following the Wiener approach (Wiener, 1964; Rieke, 2001; Figures 4A&B). Next, we convolved the estimated filters with their corresponding skewed stimuli to obtain the “effective” stimuli perceived by cone photoreceptors. Then we calculated the skews of the “effective” stimuli and plotted them against the skews of the original light stimuli in Figure 3E, where the grey line describes the situation where the “effective” skew is equal to the original skew. Data points for the positively skewed light stimuli are lower than the grey line and higher for the negatively skewed light stimuli. Consequently, temporal filtering reduced the “effective” skewness perceived by the cones.

**Figure 4.**
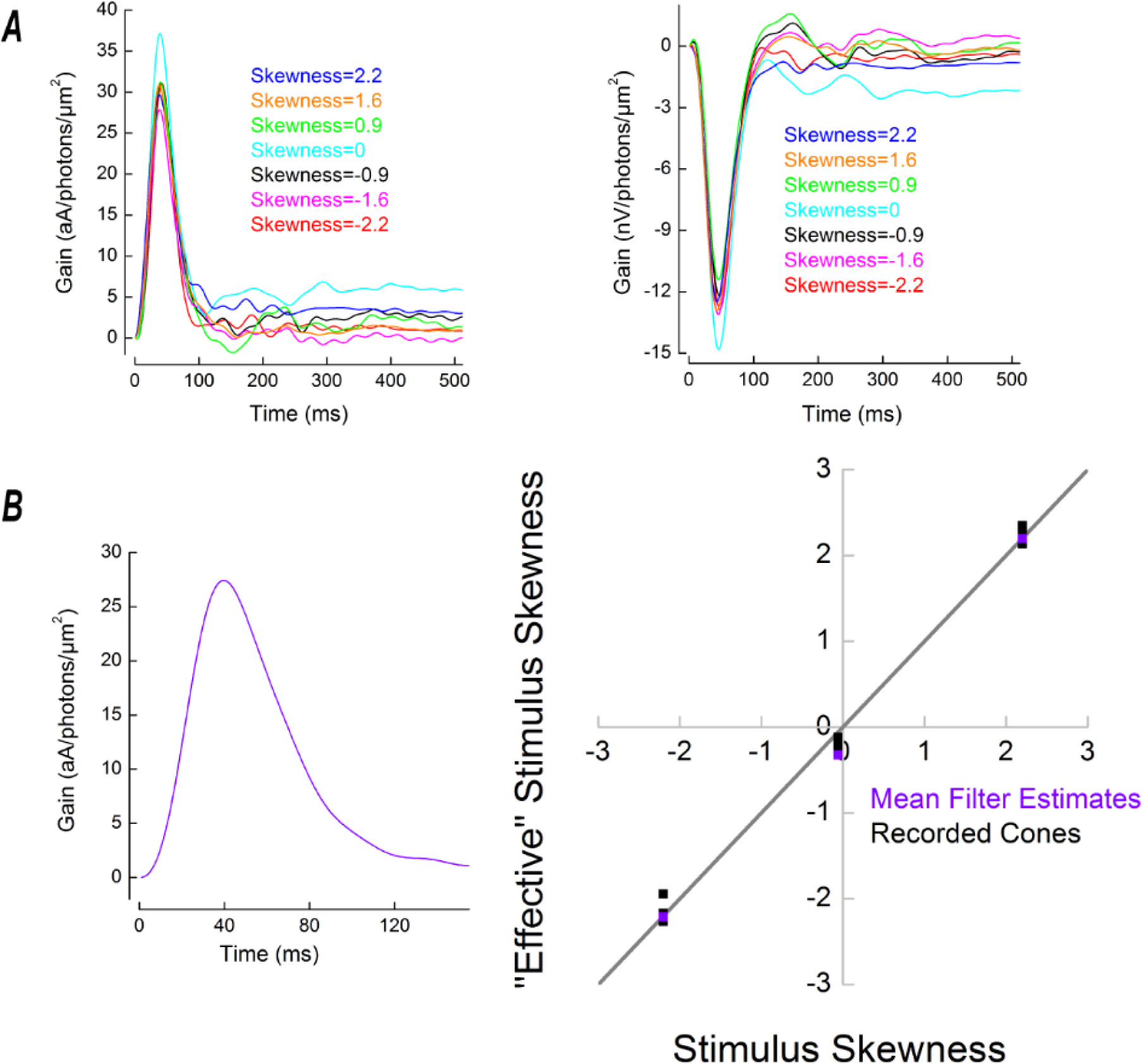
A. Representative examples of the cone impulse response functions obtained using the photocurrent (left) and voltage responses (right) to Skew Stimulus set #1. **B.** Left. The mean cone impulse response function obtained as the averaged photocurrent impulse response functions to all of the stimuli in Skew stimulus set#1 in all of the recorded cells (n=8) (Figure 4A left). This mean photocurrent impulse response function was used, via convolution, to identify segments of the NTSCI where the skewness of the original and “effective” stimuli remained equal, a subsection of which formed Skew stimulus set#2. Right. Skewness of the “effective” stimuli as a function of the skewness of the original stimuli for Skew Stimulus set #2. Violet squares correspond to the “effective” skewness obtained by the convolution of the light stimuli with the mean impulse response function shown on the left. Black squares depict the “effective” skewness ‘perceived by each cone under Skew Stimulus set #2 conditions. This was estimated by convolving each Skew Stimulus set #2 stimulus with the cone’s impulse response function obtained for corresponding stimulus. The grey line describes the condition where temporal filtering does not affect stimulus skewness. Since all squares are aligned with the grey line, the “effective” skewness is approximately equal to the original light stimulus skewness. Hence, for Skew Stimulus set #2 cone temporal filtering does not change the skewness delivered by the stimuli.

### Asymmetry in the responses to “effective” stimuli

How do goldfish cones process these “effective” stimuli? Figure 3E shows that linear temporal filtering reduces “effective” skewness and decreases the dynamic range over which responses to skewed stimuli were measured by almost 30% (from ±2.2 to ±1.6). Therefore, we first completed our data set by recording the cone’s voltage responses to stimuli with “effective” skews of ±2.2 (Figures 4C&5A). Next, we plotted the skews of the responses against the “effective” stimulus skews (Figure 5B) and found that goldfish cones decrease the magnitude of skewness when the stimuli are skewed positively and increase the magnitude of skewness when the stimuli are skewed negatively.

**Figure 5.**
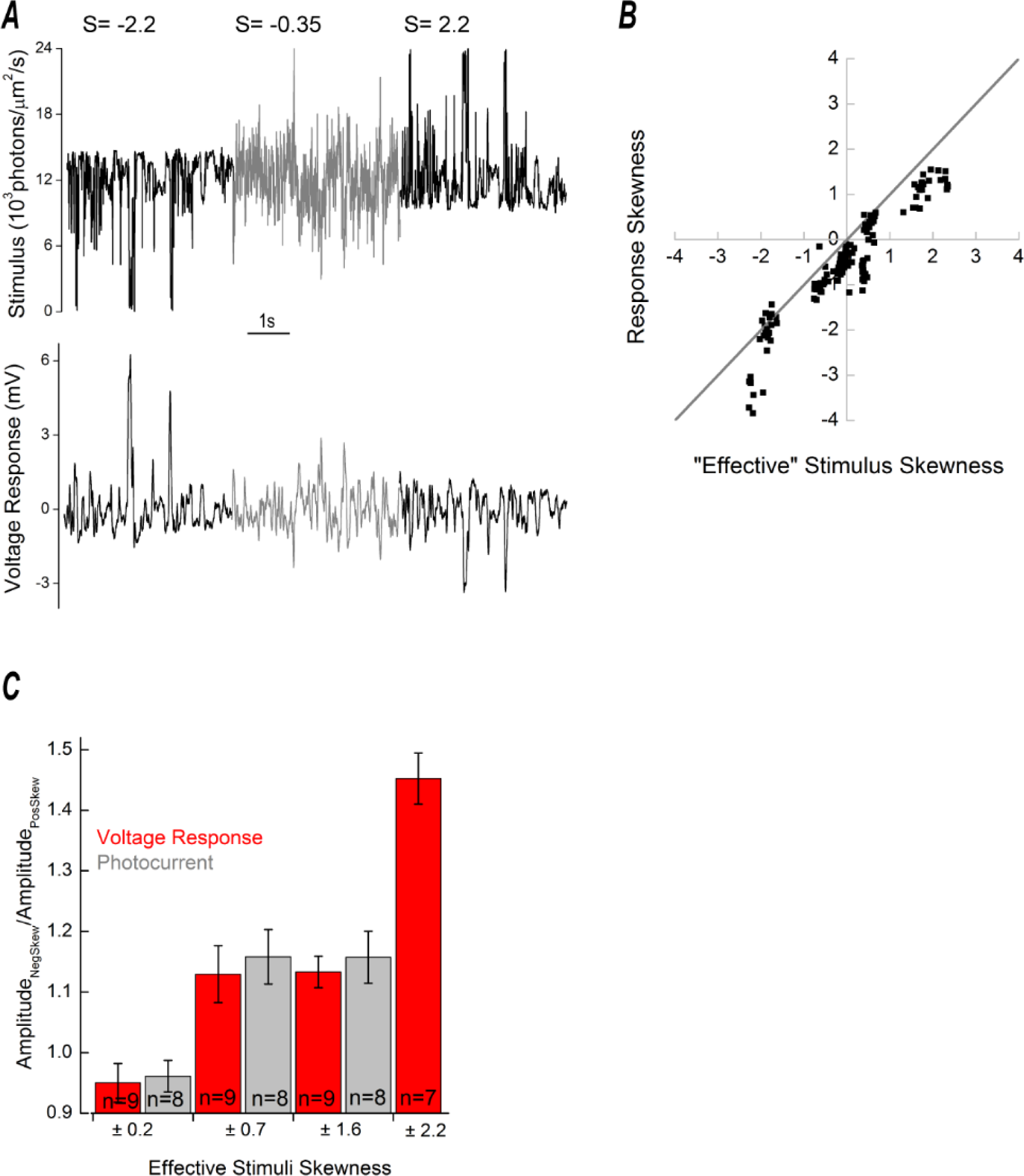
Asymmetries in cone responses to positively and negatively skewed stimuli. **A**. Top. Depiction of Skew Stimulus set #2. The key property of this stimulus set is that the light stimulus skewness is not affected by the temporal filtering properties of cones (Figure 3B). Bottom. An example of a recorded voltage response to Skew Stimulus set #2. **B**. Skews voltage and photocurrent responses against the “effective” stimuli skews. This figure illustrates the relation between “effective” stimulus skew (see Figure 3E, Figure 4B) and the skewness for the photocurrent (n=8) and voltage responses (n=9) of cones stimulated with Skew Stimulus set #1, and the voltage responses of cones (n=7) recorded with Skew Stimulus set #2. This data indicates the cone response skewness differ from the skewness of the “effective” stimulus. Cone responses are less skewed than the stimulus when it delivers higher levels of positive “effective” skew. The opposite occurs when the stimulus delivers higher levels of negative “effective” skew, the cone responses are more skewed than the stimulus. The grey line depicts the situation where the cone response and “effective” stimulus are equally skewed. For illustrative convenience, the voltage responses skews were multiplied by -1. **C**. Differences in cone response amplitudes (photocurrent – grey, voltage response – red) to negatively and positively skewed stimuli for each “effective” skew stimuli pair. When the “effective” skew magnitude was low (±0.2) the photocurrent or voltage responses amplitudes were unaffected by the skew direction. However, at higher “effective” skew-magnitudes cone response amplitudes to negatively skewed stimuli were larger than for positively skewed stimuli (photocurrent difference: ± 0.7, 15.8% ± 4.52% p=0.01; ± 1.6, 15.7 ± 4.33% p=0.0083; voltage response difference: ± 0.7, 13.0 ± 4.67% p=0.024; ± 1.6, 13.3 ± 2.64% p=0.0.00097; ± 2.2, 45.2 ± 4.20 % p=0.00004). Changes in response amplitude were assessed as the ratio of standard deviations of a cell’s response to corresponding negatively and positivity skewed stimuli. The “effective” skew values shown were estimated by the mean impulse response (Figure 7), obtained using the impulse response functions of all cells under all stimuli conditions. Data shown as mean ± SEM.

Asymmetry in the processing of negatively and positively skewed stimuli is also reflected in the amplitudes of the corresponding responses: standard deviations of the responses to negatively skewed stimuli were up to 50% larger than the standard deviations of the responses to positively skewed stimuli (Figure 5C).

### Asymmetry in the cone’s output

To be perceived by the downstream neurons, asymmetries in the cone’s responses to positively and negatively skewed stimuli (Figures 5B&C) should be reflected in the synaptic release. In photoreceptors, glutamate release is directly proportional to the calcium current (Schmitz & Witkovsky, 1997; Thoreson *et al*., 2004). Consequently, one can estimate changes in cone glutamate release by recording its calcium current (I_Ca_).

We measured skew-dependent modulation of I_Ca_ by using recorded voltage responses to light stimuli with “effective” skews of ±2.2 and -0.35 as the command voltages at three different potentials (-30, -40, -50 mV) along the I_Ca_ activation curve (Figure 6A). To isolate I_Ca_ responses, we blocked all other active conductance and subtracted the leak current. We then plotted the skews of the I_Ca_ signals against the skews of the voltage responses (Figure 6B). Note, that since depolarization produces an inward I_Ca,_ the skews of the voltage response and I_Ca_ response have opposite signs.

**Figure 6.**
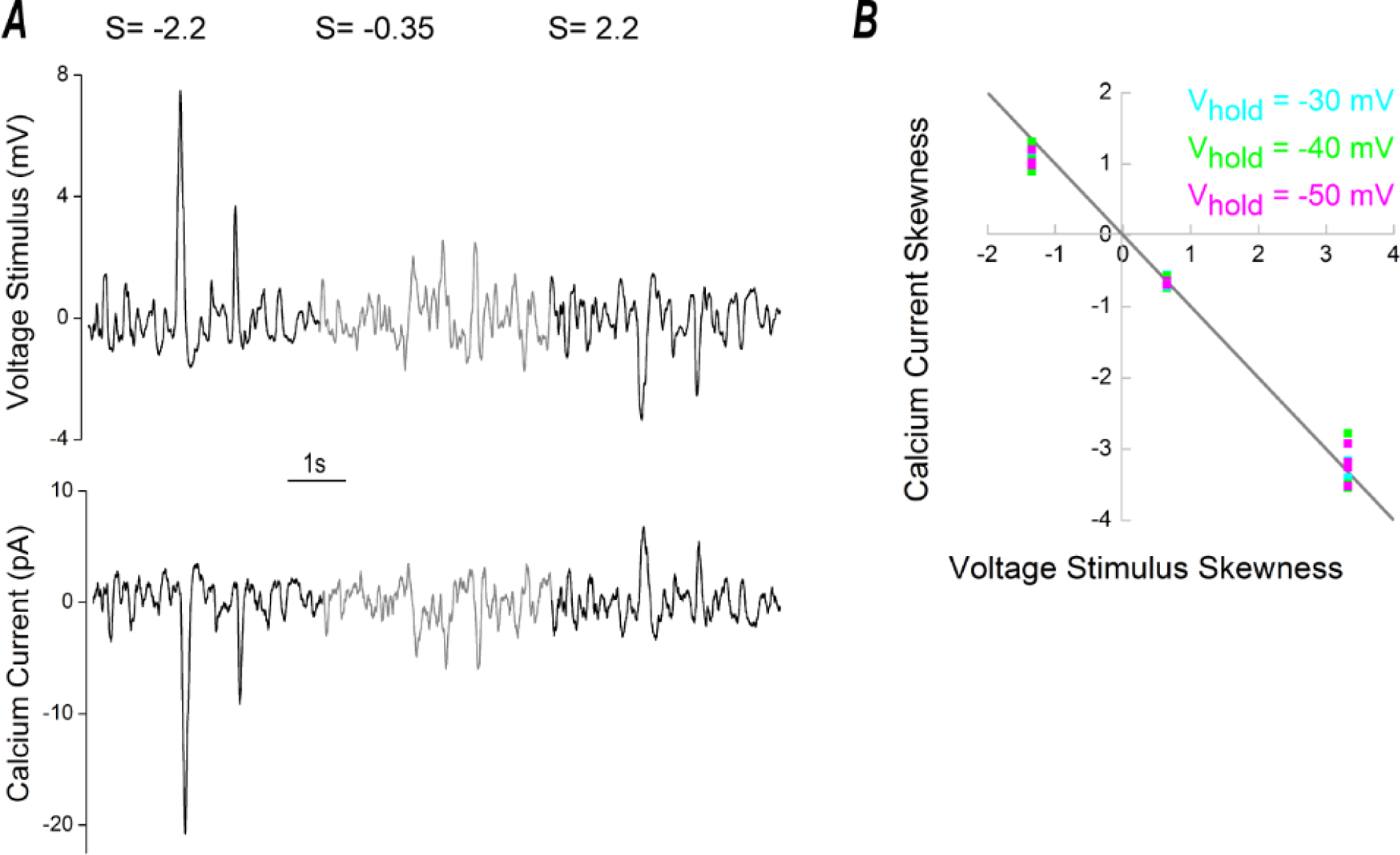
Asymmetrical processing of skewed stimuli at the cone ICa level. **A.** Top panel depicts the mean cone voltage response to Skew Stimulus set #2, which was used as the stimulus for ICa measurements. An example ICa measurement, when using this stimulus, is shown in the bottom panel. **B.** Skewness of ICa as a function of stimulus skewness. The measurements of ICa were performed at three different potentials along its activation curve: -30mV (cyan), -40mV (green) and -50mV (magenta). Note that as a decrease in voltage causes an increase in ICa, skews for the stimulus and response have opposite signs. The grey line denotes the situation where ICa and its stimulus are equally skewed. Regardless of the holding potential, data points are aligned with grey line, indicating that the cone’s ICa maintains any skewness present in its voltage response. Since cone-photoreceptor glutamate-release is directly proportional to ICa (Schmitz & Witkovsky, 1997; Thoreson *et al*., 2004), it is highly likely cone output retains any ICa skewness.

Regardless of the clamping potential, all the data points in Figure 6B approximately fall on the grey line, indicating that the skewness of the cone’s signal is largely unaffected by the transformation from membrane potential to the I_Ca_. Consistent with this, the amplitudes of I_Ca_ during the +2.2 and -2.2 skew conditions differed to the same degree (from 50 ± 1.2% to 54 ± 1% depending on the holding potential) as those of the voltage responses (Figure 5C). Thus, for negatively and positively skewed stimuli, the asymmetries present at the cone’s earlier processing stages are preserved and even somewhat enhanced in the cone’s output.

### Asymmetrical gain of the cone photoreceptors

What type of non-linear gain leads to the skew-dependent changes in the amplitude of the cone responses? To determine how the voltage response amplitude depends on the Weber contrast step we first converted the “effective” stimuli intensities into Weber contrast steps (Figure 7). Next, we plotted baseline subtracted mean voltage responses (Figure 7) as a function of the “effective” Weber contrast steps (Figure 8A). Figure 8A shows that cone responses are larger for Weber contrast steps below -0.4 than they are for Weber contrast steps above 0.4. Hence, the response gain of cones is greater for high negative, than for high positive, contrasts.

**Figure 7.**
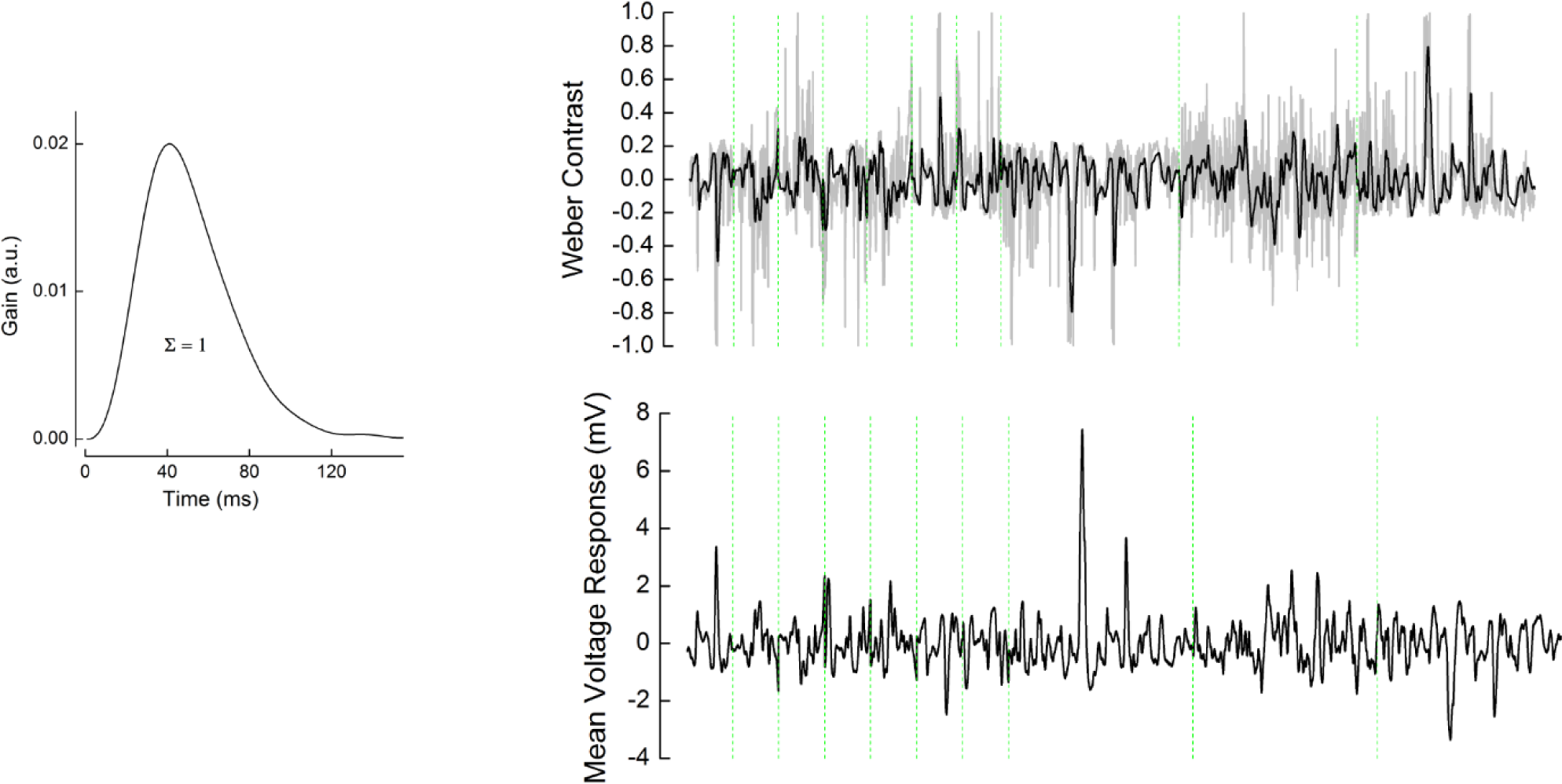
“Effective” Weber contrast steps. Left. Mean impulse response function used to obtain “effective” Weber contrast. The mean impulse response was estimated by averaging the individual voltage-responses impulse-response functions of all cells for all stimuli (n=16) (Figure 4A right, Figure 8B). This mean was scaled such that the integral under its curve was 1. Right. Upper. Grey line: The original light stimuli of Skew Stimulus set #1 and set#2 converted to Weber contrast steps. Black line: The “effective” Weber contrast steps obtained by the convolution of the original Weber contrast steps with the mean impulse response function shown on the left. Bottom. The averaged cone voltage response to each stimuli. For both upper and lower panels, the green dashed lines separate the different stimuli stretches during which the effective skews were (from left to the right): -1.6, - 0.2, 0.2, -0.7, 0.06, 1.6, 0.7, -2.2, -0.35, 2.2.

**Figure 8.**
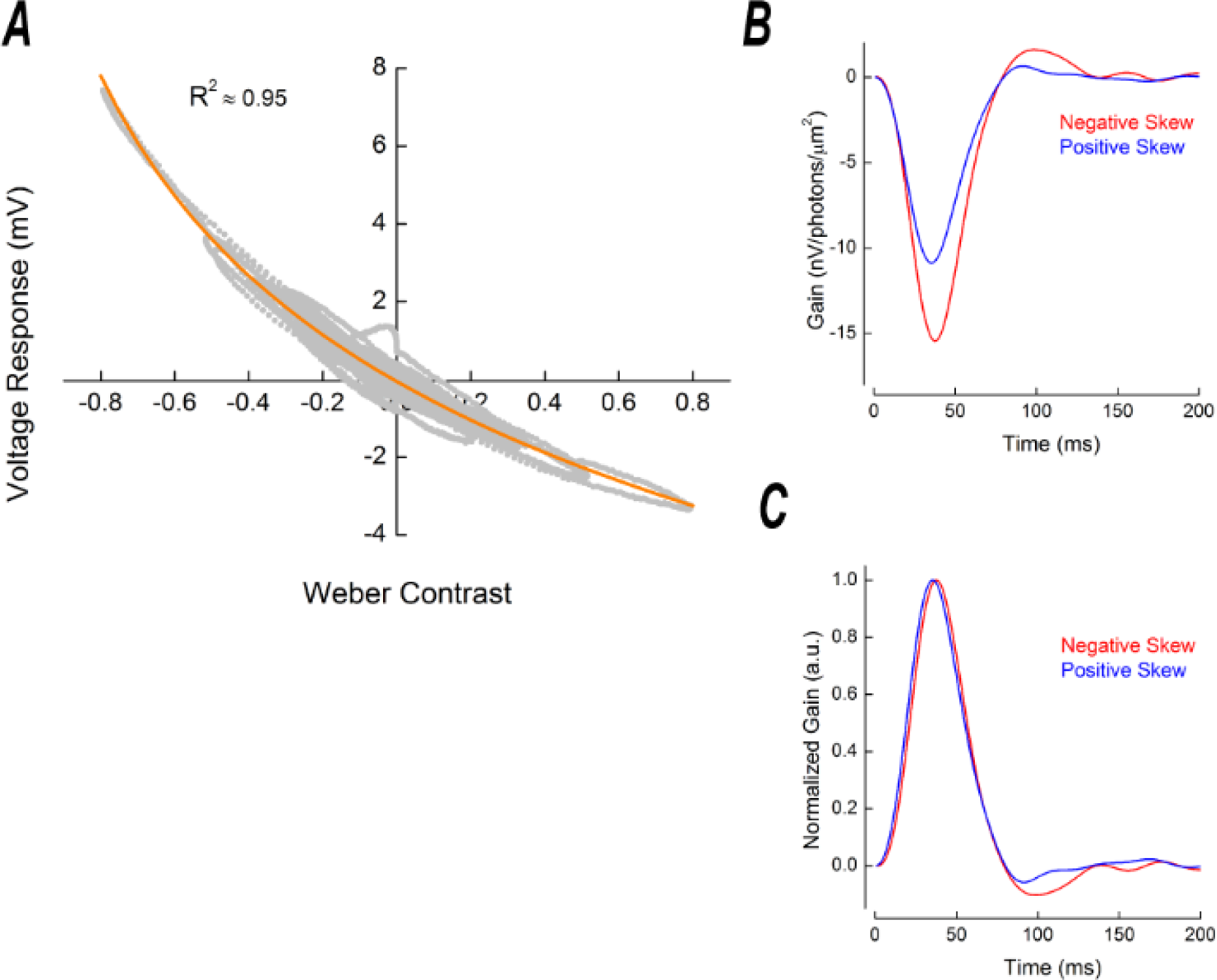
The cone signal transfer properties. **A.** Asymmetric gain of cone photoreceptors. To estimate cone photoreceptor gain, we plotted the voltage response of cones as a function of “effective” Weber contrast. This relationship was well described (orange line, adjusted R^2^ = 0.95) by the Weber contrast power function given in equation (3) and clearly indicates that cone gain is higher for negative contrasts than for the positive contrasts. The differences become prominent from Weber contrast steps around ±0.4 and are especially vivid for contrast steps beyond ±0.6. “Effective” Weber contrast was obtained by converting the light intensities into Weber contrast steps with equation (2), which were then convoluted with the mean cone impulse response function scaled such that integral under its curve yielded 1 (see methods, Figure 7). The voltage response shown is the baseline-subtracted average of all cells for each stimulus conditions. In total, there are 19000 data points in this figure. **B.** Stimulus skewness changes the shape of the cone’s impulse response function. The figure depicts two representative examples of a cone’s impulse response function in conditions with -2.2 (red) and +2.2 (blue) “effective’’ stimulus skewness. **C.** The cone impulse response functions shown in **B**, normalized by the amplitude of their initial lobe. On average, impulse response functions peaked 3.6 ms, or 9 ± 1.0 %, later for negatively skewed stimulus than for the positively skewed condition (p=0.0001, n=7) whereas the impulse response function FWHM (Δ = 1± 1.35 %, p=0.37, n=7) and the cone integration time (Δ =1.1 ± 1.61 % ,p = 0.51, n=7) were unchanged.

What kind of input-output relation supports the asymmetric gain of cones? We fitted the relation between voltage responses and “effective” Weber contrast steps with equation (3) (Figure 8A orange) and found that the cone voltage response is proportional to Weber contrast step with an exponent of -0.1251 (R^2^≈ 0.95, 95% confidence interval ± 0.0959). This dependence is close to the one determined in the theoretical study by Van Hateren and Snippe (2006), who suggested that voltage responses of the vertebrate cone is proportional to Weber contrast with an exponent of -0.12.

### Stimulus skewness affects the shape of the cone impulse response function

The processing time of cones is inversely proportional to light intensity such that responses to steps of strong positive contrast peak earlier than responses to strong negative contrasts (Nikonov *et al*., 2000; Lee *et al*., 2003; van Hateren, 2005; Van Hateren & Snippe, 2006; Angueyra *et al*., 2021). Therefore, one might expect that such a dependency would lead to a difference in the time courses of the responses to positively and negatively skewed stimuli.

To test whether stimulus skewness has an effect on the cone kinetics, we compared impulse response functions derived from the voltage responses to the -2.2 and the +2.2 “effective” skew stimuli (Figure 8B). To better visualize the differences in kinetics we normalized these impulse response functions by the amplitude of their initial lobe (Figure 8C). Interestingly, while on average the cone impulse response functions peaked 3.6 ms (or 9 ± 1.0 %) later for the negatively skewed stimulus than for the positively skewed stimulus (Figure 8C, p=0.0001, n=7), there were no statistically significant differences neither in its full width at the half maximum (FWHM) (Δ = 1.0 ± 1.35 %; p=0.37; n=7), nor in the cone’s integration time (Δ=1.1 ± 1.61 %; p=0.51; n=7). For the FWHM and integration time to remain unchanged while the time to peak shifts suggests that during negatively skewed stimuli the reduced rise time of the impulse response function’s initial flank is largely offset by the peak falling back to baseline at a faster rate.

## Discussion

We studied responses of cone photoreceptors to differently skewed stimuli and found cone response amplitudes to negatively skewed stimuli are up to 50% greater than to positively skewed stimuli (Figures 2, 5A&C, 6A, 7). This amplitude difference originates from the asymmetrical weighting of positive and negative contrasts by the phototransduction cascade. Its gain is inversely proportional to Weber contrast steps raised to the power of -0.125 (Van Hateren and Snippe, 2006; Figure 8A) and may serve as the basis for the Blackshot mechanism proposed by Chubb et al. (1994, 2004). Additionally, we observed stimulus skewness changes the cone’s impulse response function shape. For the normalized impulse response function, the rising flank was faster and the falling flank slower for positively compared to negatively skewed stimuli (Figure 8C).

### The Blackshot mechanism

Psychophysical studies reported that humans can discriminate visual stimuli based on skewness (Chubb *et al*., 1994, 2004; Graham *et al*., 2016). Our results suggest that this discrimination starts as early as the phototransduction cascade. Chubb et al. (1994, 2004) described the sensitivity to skewness with the so-called Blackshot mechanism, which has a disproportionally strong response to high negative contrasts. Our data indicates that differences in the response to positively and negatively skewed stimuli originates in the phototransduction’s asymmetric gain function, which leads to higher response amplitudes to negative contrasts than to positive contrasts (Figure 5C, Figure 8A). Moreover, in full accordance with psychophysical studies, the difference in cone response amplitudes to positive and negative contrasts becomes more prominent with larger contrast steps (Figure 8A). For the ±1.6 “effective” skew stimuli pair, where the maximal “effective” Weber contrast step was 0.5, the difference in the response was about 15% while for the ±2.2 “effective” skew pair, where the maximal “effective” Weber contrast step was 0.8, the difference was almost 50% (Figure 5C).

Asymmetries in responses to positive and negative contrasts are reported throughout the entire visual system in various species (Laughlin, 1981; Van Hateren, 1997; Lee *et al*., 2003; Zaghloul *et al*., 2003; Jin *et al*., 2008; Yeh *et al*., 2009; Endeman & Kamermans, 2010; Baden *et al*., 2013; Kremkow *et al*., 2014; Cooper & Norcia, 2015), including the human visual cortex (Zemon *et al*., 1988; Kremkow *et al*., 2014). Although it was shown that differences in response amplitudes to positive and negative contrasts are additionally amplified by the visual cortex (Kremkow *et al*., 2014), our data clearly indicates that the primary origin of this asymmetry is within the cone’s phototransduction cascade.

The phototransduction’s asymmetric gain function enables cones to efficiently encode the entire range of contrasts present in natural scenes. Photoreceptors encode changes in their input with a graded output. Information theory states that such a system encodes a signal efficiently only when the statistical distribution of its output is Gaussian, which implies a symmetrical engagement of the system’s dynamic range (Shannon, 1948; Van Hateren, 1997). On the other hand, from a given mean, the light intensity cannot decrease by more than 100%, but can easily increase by many orders of magnitude. This means that the dynamic range of positive contrasts is wider than that of negative. Thus, although some visual scenes can be skewed negatively (Tkačik *et al*., 2014), the total distribution of contrasts at any given intensity is skewed positively with negative contrasts being smaller in amplitude, but more frequent than positive contrasts (Laughlin, 1983; Ruderman, 1994; Van Hateren, 1997; Ruderman *et al*., 1998; Cooper & Norcia, 2015). Consequently, to encode signals efficiently and to provide symmetrical outputs, cones compensate for this asymmetry in their input by weighting high positive contrasts with lower gain (Figure 8A), such that when stimulated with the entire range of contrasts in natural scenes, cones provide a Gaussian output (Laughlin, 1983; Van Hateren, 1997; Endeman & Kamermans, 2010). We therefore suggest that the Blackshot mechanism is simply a consequence of the more fundamental necessity to efficiently encode the range of contrasts present in natural scenes.

It is also important to note that although when it comes to dealing with wide dynamic ranges of light intensities the non-linear gain of photoreceptors is well-acknowledged, its influence on perception is often ignored such that responses of the downstream neurons are often modelled with an implicit assumption of the photoreceptor linearity (Chander & Chichilnisky, 2001; Kim & Rieke, 2001; Rieke, 2001; Carandini *et al*., 2005; Mante *et al*., 2005; Manookin & Demb, 2006; Ozuysal & Baccus, 2012; Pitkow & Meister, 2012; Karamanlis & Gollisch, 2021; Schreyer & Gollisch, 2021). However, there are a number of occasions where this non-linearity is important to account for perceptual features (Kremkow *et al*., 2014; Angueyra *et al*., 2021). For instance, higher spatial resolution for darker than for brighter patches is a consequence of the cone photoreceptor non-linear gain (Kremkow *et al*., 2014). Our data presents another example of the far-reaching consequences of the early visual non-linearity and highlights importance of accounting for the photoreceptor non-linearity when studying the visual system.

### Shape of the impulse response function

We found the cone’s impulse response function peaks ≈ 3.6 ms later for negatively skewed stimuli whereas the cone’s integration time is unaffected by stimulus skewness (Figure 8B&C). The rising and falling flanks of the cone’s impulse response are governed by different biophysical mechanisms. The former is heavily influenced by the phosphodiesterase (PDE) hydrolysis of cGMP. The time constant of this process is inversely proportional to the light intensity. Hence, it decreases upon positive, and increases upon negative contrast steps (Nikonov *et al*., 2000; van Hateren, 2005; Endeman & Kamermans, 2010). The terminating flank is largely regulated by the guanylyl cyclase (GC) mediated production of cGMP which is modulated by the Ca^2+^ influx through the CNG channels. The time constant of this process is not light-dependent but its gain is inversely proportional to the 4th power of light intensity (Burns et al., 2002; van Hateren, 2005; van Hateren and Snippe, 2007). The interplay between these two underlying processes is thought to account for the cone’s impulse response function shape, the asymmetric rising and falling response phases to sinusoidal stimuli, and for light adaptation to decreases in light intensity being slower than for light intensity increases (Baylor & Hodgkin, 1973; Lankheet *et al*., 1991; Nikonov *et al*., 2000; Lee *et al*., 2003; van Hateren, 2005; Endeman & Kamermans, 2010; Angueyra *et al*., 2021).

For our stimuli, the PDE hydrolysis of cGMP time-constant will have been shorter during positively skewed stimuli as all large changes in light intensity were associated with positive contrasts. This in turn manifest as a faster rate of change in the impulse response functions’ initial flank and hence an earlier time to peak. For negatively skewed stimuli, as all large changes in light intensity were associated with negative contrast, GC mediated production of cGMP was pronounced. The highly non-linear light-dependent gain function of this process, and the ensuing Ca^2+^ influx when cones depolarized, increased the rate the cone CNG channels reopened. This resulted in an increased decay rate for the terminating flank of the impulse response function. Hence, the initial flank’s faster onset rate during the positively skewed stimulus is largely offset by the terminating flank’s faster decay rate during the negatively skewed stimulus. Consequently, the cone impulse response function integration time to positive and negative skewed stimuli does not differ while it’s time to peak does.

### Relation to previous studies using skewed stimuli

Why did previous studies find stimulus skewness had little to no effect on RGC (Tkačik *et al*., 2014) and LGN neurons (Bonin *et al*., 2006)? We suggest a methodological factor. In both studies, a large proportion of the stimulus power spectrum was outside the cone’s temporal frequency bandwidth. Indeed, Bonin et al. (2006) used white-noise stimuli bandlimited to 124 Hz to study cat LGN neurons responses, while the cat visual system barely responds to frequencies above 32 Hz (Shapley & Victor, 1978; Mante *et al*., 2005). Similarly, Tkacik et al. (2014) studied salamander RGC responses with white-noise bandlimited to 30 Hz, whereas the salamander retina hardly reacts to frequencies above 10 Hz (Kim & Rieke, 2001). Thus, in both these studies a large part of their stimuli were ‘filtered out’ and the remaining “effective” stimuli were only able to elicit marginal skew dependent effects.

To illustrate this point, we estimated the “effective” stimuli delivered by Bonin et al. (2006) and Tkacik et al. (2014) using Van Hateren’s cone photoreceptor model (van Hateren & Snippe, 2007). Simulations of the cat cone indicate that while a wide range of “effective” Weber contrasts were present (Figure 9A) the “effective” skewness was approximately half that of the original stimuli employed, reducing from a range of ±0.4 to approximately ±0.2 (Figure 9B). Hence, both stimuli delivered largely similar distributions of “effective” Weber contrasts. As a result, the simulated voltage responses to the positive and negatively skewed stimuli only differed in amplitude by approximately 4% (Figure 9 C, D), which is within the range of standard error estimates for the amplitude differences we find here (Figure 5C).

**Figure 9.**
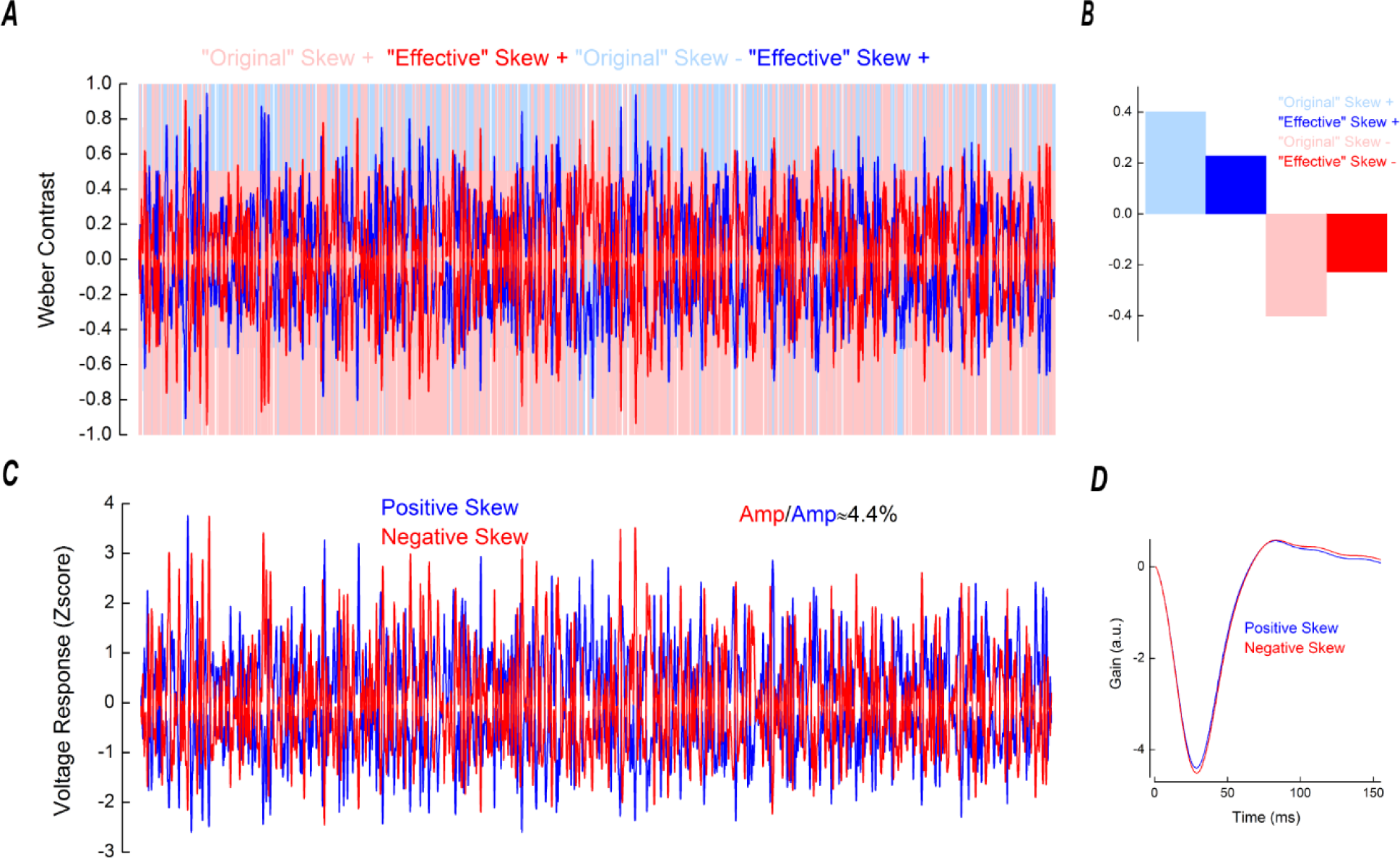
Simulated responses of cat cone photoreceptors. **A.** Light blue and light red lines depict the positively and negatively skewed stimuli used by Bonin et al. (2006) and here for the simulation. Blue and red lines depict the “effective” positively and negatively skewed stimuli, obtained by the convolution of the “original” stimuli with the impulse response functions shown in **D**. **B.** Comparison of the skews of the “original” and “effective” positively (blue) and “negatively” (red) skewed stimuli. Due to the temporal filtering the “effective” stimuli are almost symmetrical around the mean, their skew range having decreased from ±0.4 to ≈ ±0.2. **C.** Simulated cone voltage responses to the positively (blue) and negatively (red) skewed stimuli. The response amplitude to the negatively skewed stimulus was 4% higher than the response amplitude to the positively skewed stimulus. The simulated voltage response amplitudes were quantified by their standard deviations, as for Figure 5C. Parameters for the simulation are listed in the Table 1. **D.** Impulse response functions derived from the simulated cat cone voltage responses. These impulse response functions were used to estimate the “effective” stimuli shown in **A.**

Simulations of the salamander cone reveal a different situation that none the less leads to the same outcome. The “effective” skewness range remained relatively large despite being less than half that of the original stimuli employed (±0.8 vs ±2, Figure 10B), but the range of “effective” contrasts reduced to just ±0.2 Weber unit (Figure 10A). Over this limited range of contrasts the cone photoreceptor gain is mostly symmetrical (Figure 8A) and as such the voltage response amplitude to the negatively skewed stimulus was only 2% higher than for the positively skewed stimulus (Figure 10C&D). Hence, even though the stimuli had substantially different distributions of “effective” Weber contrasts, the range of contrast values they delivered were too narrow to generate a notable effect.

**Figure 10.**
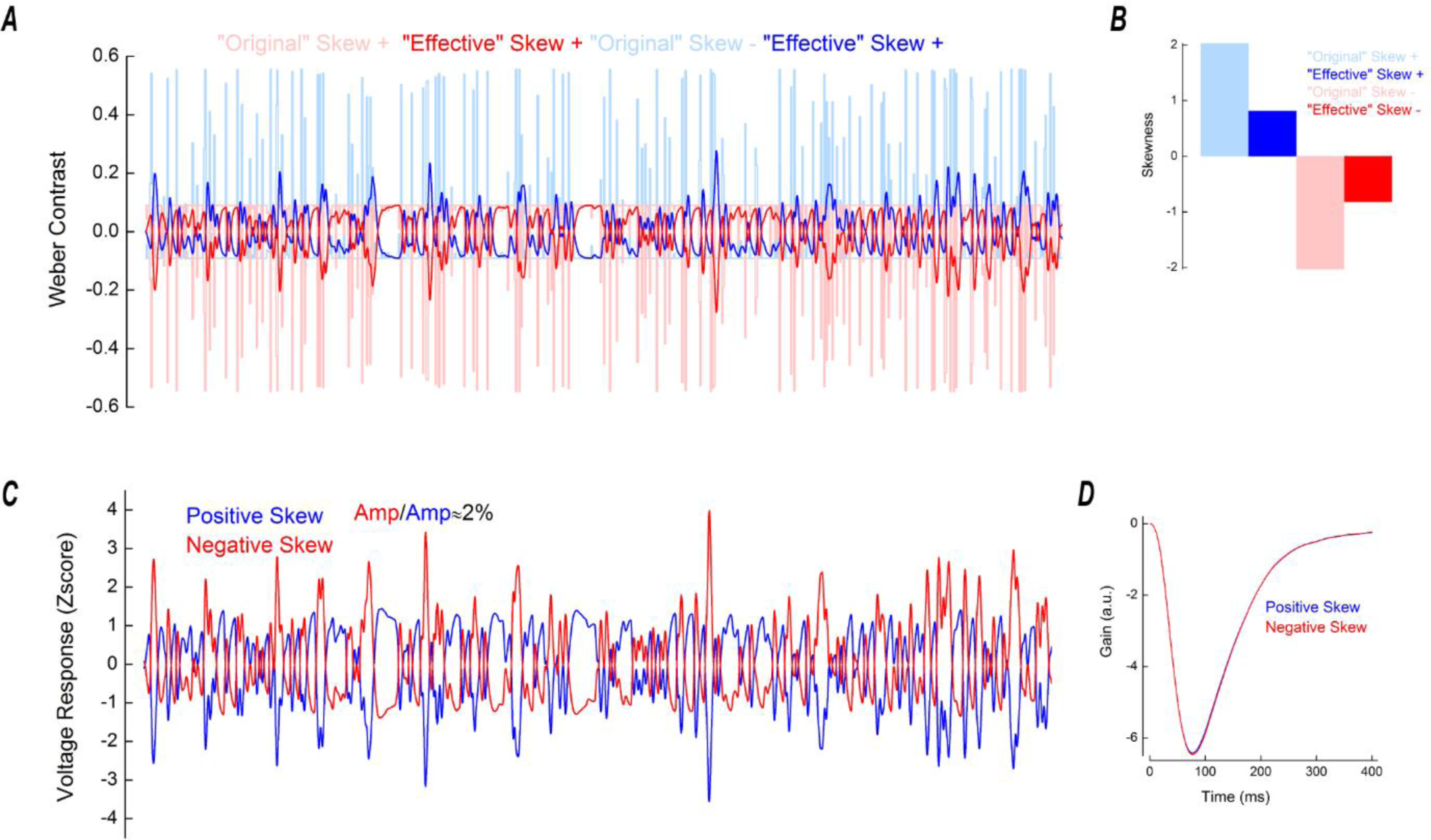
Simulated responses of salamander cone photoreceptors. **A.** Light blue and light red lines depict the positively and negatively skewed stimuli used in the simulation. These stimuli had the same properties as those used by Tkacik et al.(2014). Blue and red lines depict the “effective” positively and negatively skewed stimuli, obtained by the convolution of the “original” stimuli with the impulse response functions shown in **D**. The temporal filtering leads to small Weber contrast ranges of the “effective” stimuli. **B.** Comparison of the skews of the “original” and “effective” positively (blue) and “negatively” (red) skewed stimuli. The salamander cone’s temporal filtering reduced the stimulus skewness from ±2 to ±0.8. **C.** Simulated cone voltage responses to the positively (blue) and negatively (red) skewed stimuli. The response amplitude to the negatively skewed stimulus was 2% higher than the response amplitude to the positively skewed stimulus. The simulated voltage response amplitudes were quantified by their standard deviations, as for Figure 5C. Parameters for the simulation are listed in the Table 1. **D.** Impulse response functions derived from the simulated salamander cone voltage responses. These impulse response functions were used to estimate the “effective” stimuli shown in **A.**

To conclude, our results show that to study visual processes under varying skewness conditions, the stimuli must be able to deliver sufficient levels of “effective” skewness over sufficient ranges of “effective” Weber contrasts.

